# RAD sequencing and a hybrid Antarctic fur seal genome assembly reveal rapidly decaying linkage disequilibrium, global population structure and evidence for inbreeding

**DOI:** 10.1101/269167

**Authors:** Emily Humble, Kanchon K Dasmahapatra, Alvaro Martinez-Barrio, Inês Gregório, Jaume Forcada, Ann-Christin Polikeit, Simon D Goldsworthy, Michael E Goebel, Joern Kalinowski, Jochen Wolf, Joseph I Hoffman

## Abstract

Recent advances in high throughput sequencing have transformed the study of wild organisms by facilitating the generation of high quality genome assemblies and dense genetic marker datasets. These resources have the potential to significantly advance our understanding of diverse phenomena at the level of species, populations and individuals, ranging from patterns of synteny through rates of linkage disequilibrium (LD) decay and population structure to individual inbreeding. Consequently, we used PacBio sequencing to refine an existing Antarctic fur seal (*Arctocephalus gazella*) genome assembly and genotyped 83 individuals from six populations using restriction site associated DNA (RAD) sequencing. The resulting hybrid genome comprised 6,169 scaffolds with an N50 of 6.21 Mb and provided clear evidence for the conservation of large chromosomal segments between the fur seal and dog (*Canis lupus familiaris*). Focusing on the most extensively sampled population of South Georgia, we found that LD decayed rapidly, reaching the background level of *r*^2^ = 0.09 by around 26 kb, consistent with other vertebrates but at odds with the notion that fur seals experienced a strong historical bottleneck. We also found evidence for population structuring, with four main Antarctic island groups being resolved. Finally, appreciable variance in individual inbreeding could be detected, reflecting the strong polygyny and site fidelity of the species. Overall, our study contributes important resources for future genomic studies of fur seals and other pinnipeds while also providing a clear example of how high throughput sequencing can generate diverse biological insights at multiple levels of organisation.

## INTRODUCTION

Advances in short read sequencing technologies, in particular Illumina sequencing, have made it possible to generate genome assemblies as well as dense genetic marker datasets for practically any organism (Ekblom and Galindo 2011; Ellegren 2014). However, assemblies based solely on short read data tend to be highly fragmented, even with assembly strategies that incorporate medium length insert libraries (Gnerre *et al.* 2011). Consequently, although such assemblies can be generated rapidly and cheaply, there has been growing interest in technologies that incorporate longer range information to improve scaffold length and contiguity. For example, Pacific Biosciences (PacBio) single molecule real-time (SMRT) sequencing generates read lengths in the order of several kilobases (kb) that have proven effective in gap filling, resolving complex repeats and increasing contig lengths across diverse taxa (English *et al.* 2012; Conte and Kocher 2015; Pootakham *et al.* 2017).

In parallel to these and related developments in genome sequencing technologies, reduced representation sequencing approaches such as restriction site associated DNA (RAD) sequencing (Baird *et al.* 2008; Peterson *et al.* 2012) are providing unprecedented levels of genetic resolution for population genetic and genomic studies (Morin *et al.* 2004; Stapley *et al.* 2010; Seeb *et al.* 2011). By sequencing and assembling short stretches of DNA adjacent to restriction cut sites and interrogating the resulting tags for sequence polymorphisms, RAD sequencing can facilitate the acquisition of large genome-wide distributed single nucleotide polymorphism (SNP) datasets incorporating multiple individuals.

The above approaches show great promise for studying wild populations where genomic resources are typically absent. For example, information from model organisms with well-characterised genomes can facilitate studies of their wild relatives as long as patterns of synteny between the two can be established. Knowledge of synteny can facilitate the lifting over of gene annotations, assist in gene mapping and help to elucidate the genetic basis of fitness variation by identifying genes closely linked to loci responsible for inbreeding depression (Johnston *et al.* 2011; Ekblom and Wolf 2014; Kardos *et al.* 2016).

High density SNP markers mapped to a reference genome can furthermore provide insights into processes that shape levels of variation within genomes. For example, the positional information of genomic loci can be used to characterise patterns of linkage disequilibrium (LD). LD is a central concept in population genetics because it is closely associated with factors such as effective population size (*N*_*e*_), genetic drift, historical fluctuations in population size, population structure, inbreeding and recombination (Slatkin 2008). Understanding the strength and extent of LD can aid in the inference of demographic history and has important implications for identifying genetic variants underlying key fitness traits through genome-wide association analyses or quantitative trait locus mapping (Carlson *et al.* 2004; Miller *et al.* 2015; Kardos *et al.* 2016). Nevertheless, the genomic pattern of LD has only been described in a handful of wild populations. Typically, LD decays within a few tens to hundreds of kilobases (kb) in large and unstructured populations (Poelstra *et al.* 2013; Kawakami *et al.* 2014; Vijay *et al.* 2016), but can extend for several megabases (Mb) in smaller, isolated, heavily bottlenecked and/or inbred populations, such as wolves and sheep (Hagenblad *et al.* 2009; Miller *et al.* 2011).

In addition to facilitating the characterisation of genome-wide patterns of variation, dense genomic markers can also be used to describe variation at the population and individual level, even without positional information. For example, studies are increasingly employing approaches such as RAD sequencing to obtain large datasets in order to reliably characterize genetic structure (Malenfant *et al.* 2015; Benestan *et al.* 2015; Younger *et al.* 2017) and many are uncovering patterns that had previously gone undetected (Reitzel *et al.* 2007; Ogden *et al.* 2013; Vendrami *et al.* 2017). A precise understanding of population structure is critical for the delineation of management units for conservation (Bowen *et al.* 2005) as well as for avoiding false positives in genome-wide association studies (Johnston *et al.* 2011) but can also be a used to determine contemporary and historical barriers to gene flow (McRae *et al.* 2005; Hendricks *et al.* 2017) and to elucidate patterns of extinction and recolonization (McCauley 1991).

A major topic of interest at the level of the individual is the extent to which inbreeding occurs in natural populations (Kardos *et al.* 2016) and its consequences for fitness variation and population demography (Keller and Waller 2002). Pedigree-based studies, typically of isolated island populations and often involving polygynous species, have uncovered widespread evidence of inbreeding in the wild (Marshall *et al.* 2002; Townsend and Jamieson 2013; Nietlisbach *et al.* 2017). However, the extent of inbreeding in large, continuous and free-ranging populations remains open to question. On the one hand, simulations have suggested that inbreeding will be absent from the vast majority of wild populations with the possible exception of highly polygynous and/or structured populations (Balloux *et al.* 2004). On the other hand, associations between microsatellite heterozygosity and fitness (heterozygosity fitness correlations, HFCs) have been described in hundreds of species (Chapman *et al.* 2009) and it has been argued that these are highly unlikely to arise in the absence of inbreeding (Szulkin *et al.* 2010). Due to the high sampling variance of microsatellites, there has been growing interest in the use of high density SNP data to reliably quantify inbreeding, and recent empirical and simulation studies suggest that this can be achieved with as few as 10,000 SNPs (Kardos *et al.* 2015; 2018). Consequently, with approaches like RAD sequencing, it should be possible to quantify the variation in inbreeding in arguably more representative wild populations.

The Antarctic fur seal (*Arctocephalus gazella*) is an important marine top predator that has been extensively studied for several decades, yet many fundamental aspects of its biology remain poorly understood. This highly sexually dimorphic pinniped has a circumpolar distribution and breeds on islands across the sub-Antarctic, with 95% of the population concentrated on South Georgia in the South Atlantic (Figure 1). The species was heavily exploited by 18^th^ and 19^th^ Century sealers and was thought to have gone extinct at virtually all of its contemporary breeding sites (Weddell 1825). However, in the 1930s a small breeding population was found at South Georgia (Bonner 1968; Payne 1977), which in the following decades increased to number several million individuals (Boyd 1993). While it is believed that the species former range was recolonised by emigrants from this large and rapidly expanding population (Boyd 1993; Hucke-Gaete *et al.* 2004), one would expect to find little or no population structure under such a scenario. However, a global study using mitochondrial DNA resolved two main island groups (Wynen *et al.* 2000) while microsatellites uncovered significant differences between South Georgia and the nearby South Shetland Islands (Bonin *et al.* 2013), implying that at least two relict populations must have survived sealing.

Antarctic fur seals have been intensively studied for several decades at a small breeding colony on Bird Island, South Georgia, where a scaffold walkway provides access to the animals for the collection of detailed life history and genetic data. Genetic studies have confirmed behavioural observations of strong polygyny (Bonner 1968) by showing that a handful of top males father the majority of offspring (Hoffman *et al.* 2003). Furthermore, females exhibit strong natal site fidelity, returning to within a body length of where they were born to breed (Hoffman and Forcada 2012), while adults of both sexes are highly faithful to previously held breeding locations (Hoffman *et al.* 2006). Together these behavioural traits may increase the risk of incestuous matings. In line with this, heterozygosity measured at nine microsatellites has been found to correlate with multiple fitness traits including early survival, body size and reproductive success (Hoffman *et al.* 2004; 2010; Forcada and Hoffman 2014). However, such a small panel of microsatellites cannot provide a very precise estimate of inbreeding (Slate *et al.* 2004; Balloux *et al.* 2004) and therefore high density SNP data are required to provide more detailed insights into the variance in inbreeding in the population.

Here, we used PacBio sequencing to improve an existing Antarctic fur seal genome assembly comprising 8,126 scaffolds with an N50 of 3.1 Mb (Humble *et al.* 2016). We additionally RAD sequenced 83 individuals, mainly from South Georgia but also from an additional five populations, to generate a large dataset of mapped genetic markers. The resulting data were then used to investigate synteny with the dog and, within the focal South Georgia population, to characterise the pattern of LD decay as well as variance in inbreeding. Finally, using data from both RAD sequencing and 27 microsatellites, we investigated the strength and pattern of population structure across the species range and compared the ability of the two marker types to resolve genetic differences between island groups. Our hypotheses were as follows: (i) We expected to find strong synteny between the fur seal and dog (*Canis lupus familaris*), the closest relative with an annotated, chromosome-level genome assembly; (ii) LD might be expected to decay very rapidly given that fur seals are free-ranging with large population sizes. However, the historical bottleneck could potentially have resulted in elevated levels of LD; (iii) We hypothesised that nuclear markers would detect the same two island groups as previously found with mitochondrial DNA as well as possibly resolve finer scale structuring. Furthermore, RAD sequencing should provide greater power to capture genetic differences than microsatellites; (iv) Finally, we expected to find variation in inbreeding consistent with knowledge of the species mating system as well as previous studies documenting HFCs.

## MATERIALS AND METHODS

### Hybrid genome assembly and PacBio DNA library preparation

We first used the program GapCloser v1.12 to fill gaps in the existing fur seal genome v1.02 (Humble *et al.* 2016) (NCBI SRA: BioProject PRJNA298406) based on the paired end information of the original Illumina reads. This approach closed 45,852 gaps and reduced the amount of N space in the assembly from 115,235,953 bp to 78,393,057 bp (v1.1, Table 1). Following this, we generated SMRT sequencing data from the DNA used for the original genome assembly (NCBI SRA: BioSample SAMN04159679) following the protocol described in Pendleton *et al.* (2015). First, 10 μg of pure genomic DNA was fragmented to 20 kb using the Hydroshear DNA shearing device (Digilab, Marlborough, MA) and size-selected to 9–50 kb using a Blue Pippin according to the standard Pacific Biosciences SMRT bell construction protocol. The library was then sequenced on 64 PacBio RSII SMRT cells using the P6–C4 chemistry. This yielded a total of 58 Gb (~19x) of sequencing data contained within 8,101,335 subread bases with a mean read length of 7,177 bp (median = 6,705 bp; range = 50-54,622 bp). The data have been deposited to the NCBI SRA under accession number XXXX.

Next, we used PBJelly v15.8.24 and blasr (https://github.com/PacificBiosciences/blasr) with default parameters to align the PacBio sequencing reads to the gap-closed assembly to generate a hybrid genome (v1.2). Lastly, we followed a two-step strategy to remove any indels introduced by single molecule real-time sequencing (Ross *et al.* 2013). We first used Quiver (contained in the SMRT/2.3.0 suite: GenomicConsensus v0.9.2) with the refineDinucleotideRepeats option to perform initial assembly error correction. Due to this step being computationally demanding, we ran it separately for each scaffold. Next, we mapped the original Illumina reads (Humble *et al.* 2016) to the quiver assembly (v1.3) using BWA MEM v0.7.15 (Li 2013) and used Picard tools to sort and mark duplicates. We then used PILON v1.22 (Walker *et al.* 2014) to perform the final error correction step to generate assembly v1.4. The final assembly is available at NCBI under accession number XXXX.

### Genome alignment

We aligned the fur seal scaffolds from assembly v1.4 to the dog genome (*Canis lupus familiaris* assembly version CanFam3.1, GenBank accession number GCA_000002285.2) using LAST v746 (Kiełbasa *et al.* 2011). First, the dog genome was prepared for alignment using the command lastdb. We then used lastal and last-split in combination with parallel-fastq to align the fur seal scaffolds against the dog genome. Using the program MafFilter, we then processed the resulting multiple alignment format (maf) file and estimated pairwise sequence divergence between the two species (Dutheil *et al.* 2014). Finally, we extracted alignment coordinates from the maf file using bash commands to allow subsequent visualisation with the R package RCircos (Zhang *et al.* 2013).

### Sampling and DNA extraction

Tissue samples were collected from 57 Antarctic fur seal individuals from Bird Island, South Georgia. These comprised 24 partially overlapping triads consisting of 24 pups, 16 mothers and 17 fathers. Additional samples were obtained from the main breeding colonies across the species range (Figure 1): Cape Shirreff in the South Shetlands (*n* = 6), Bouvetøya (*n* = 5), Îles Kerguelen (*n* = 5), Heard Island (*n* = 5) and Macquarie Island (*n* = 5). Skin samples were collected from the inter-digital margin of the fore-flipper using piglet ear notching pliers and stored in 20% dimethyl sulphoxide saturated with NaCl at –20°C. Skin samples from the South Shetlands were collected using a sterile 2mm biopsy punch and stored in 95% ethanol. Total genomic DNA was extracted using a standard phenol-chloroform protocol (Sambrook *et al.* 1989).

### Microsatellite genotyping

All samples were genotyped at 27 polymorphic microsatellite loci (see Supplementary table 1), previously been found to be in Hardy-Weinberg equilibrium (HWE) in the study population at South Georgia and are unlinked (Stoffel *et al.* 2015; Peters *et al.* 2016). The loci were PCR amplified in three separate multiplexed reactions (see Supplementary Table 1) using a Type It Kit (Qiagen). The following PCR profile was used for all multiplex reactions except for multiplex one: initial denaturation of 5 min at 94°C; 28 cycles of 30 sec at 94°C, 90 sec at 60°C, and 30 sec at 72°C, followed by a final extension of 30 min at 60°C. The PCR profile of multiplex one only differed from this protocol in the annealing temperature used, which was 53°C. Fluorescently labelled PCR products were then resolved by electrophoresis on an ABI 3730xl capillary sequencer and allele sizes were scored using GeneMarker v1.95. To ensure high genotype quality, all traces were manually inspected and any incorrect calls were adjusted accordingly.

**Figure 1.**
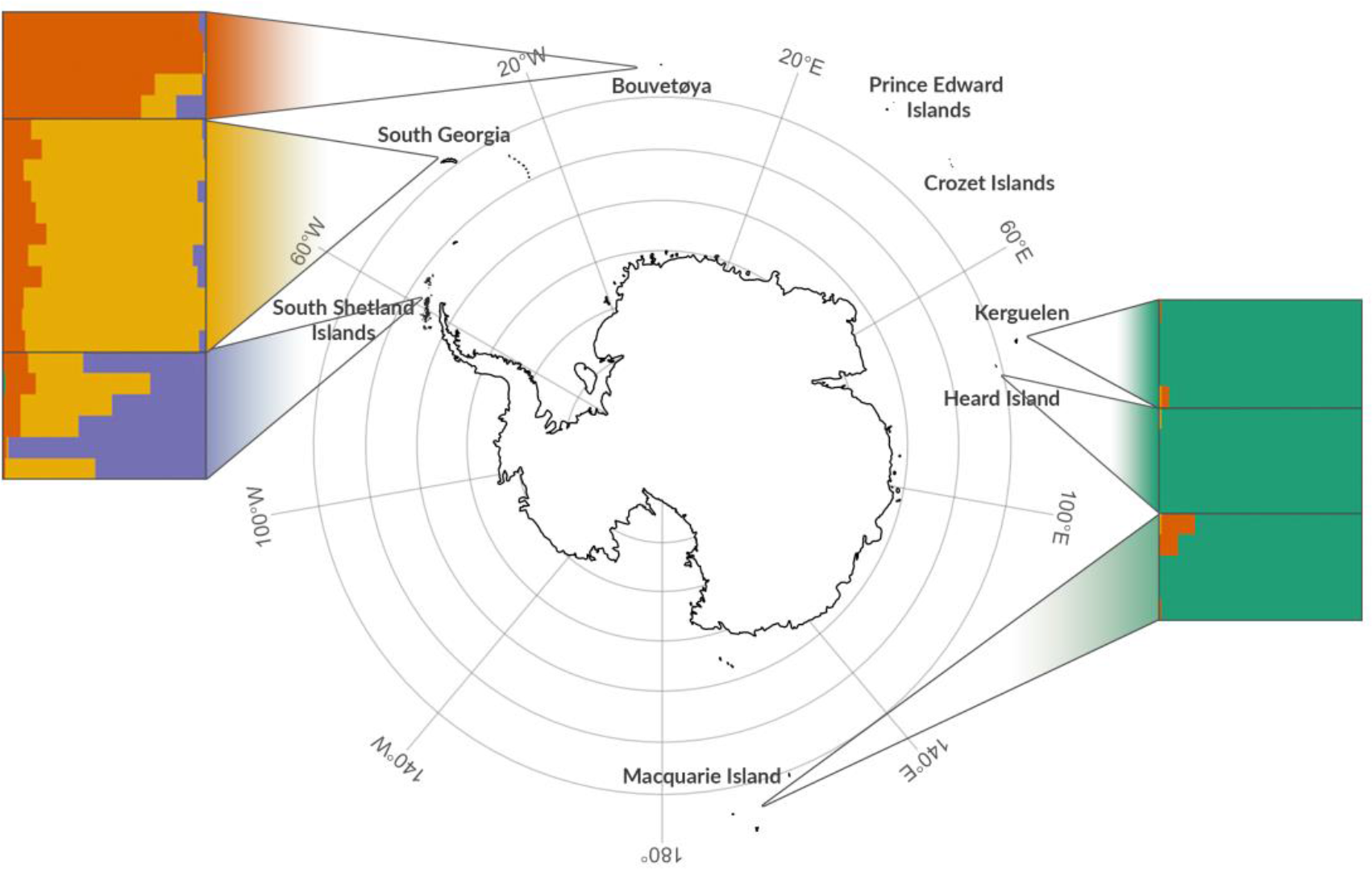
Individual assignment to genetic clusters based on STRUCTURE analysis for *K* = 4 using 28,092 SNPs. Each horizontal bar represents a different individual and the relative proportions of the different colours indicate the probabilities of belonging to each group. Individuals are separated by sampling locations as indicated on the map.

### RAD library preparation and sequencing

RAD libraries were prepared using a modified protocol from Etter *et al.* (2011) with minor modifications as described in Hoffman *et al.* (2014). Briefly, 400 ng of genomic DNA from each individual was separately digested with *SbfI* followed by the ligation of P1 adaptors with a unique 6 bp barcode for each individual in a RAD library, allowing the pooling of 16 individuals per library. Libraries were sheared with a Covaris S220 and agarose gel size-selected to 300-700 bp. Following 15-17 cycles of PCR amplification, libraries were further pooled using eight different i5 indices prior to 250 bp paired-end sequencing on two lanes of Illumina HiSeq 1500. The sequences have been deposited in the Short Read Archive (accession no. XXX).

### SNP genotyping

Read quality was assessed using FastQC v0.112 and sequences trimmed to 225 bp and demultiplexed using process_radtags in STACKS v1.41 (Catchen *et al.* 2013). We then followed GATK’s best practices workflow for variant discovery (Poplin *et al.* 2017). Briefly, individual reads were mapped to the Antarctic fur seal reference genome v1.4 using BWA MEM v0.7.10 (Li 2013) with the default parameters. Any unmapped reads were removed from the SAM alignment files using SAMtools v1.1 (Li 2011). We then used Picard Tools to sort each SAM file, add read groups and remove PCR duplicates. Prior to SNP calling, we performed indel realignment to minimize the number of mismatching bases using the RealignerTargetCreator and IndelRealigner functions in GATK v3.6. Finally, HaplotypeCaller was used to call variants separately for each individual. Genomic VCF files were then passed to GenotypeGVCFs for joint genotyping. The resulting SNP dataset was then filtered to include only biallelic SNPs using BCFtools v1.2 (Li 2011) to obtain a dataset of 677,607 SNPs genotyped in 83 individuals. Subsequently, we applied a variety of filtering steps according to the analysis being performed as shown in figure S1 and described below.

### SNP validation

To provide an indication of the quality of our SNP dataset, we attempted to validate a representative subset of loci using Sanger sequencing. First, we randomly selected 50 loci whose 70 bp flanking sequence contained no secondary SNPs and mapped uniquely to the fur seal reference genome and with initial depth of coverage and minor allele frequency (MAF) filters of 5 and 0.05 respectively. We then designed oligonucleotide primers using Primer 3 (Untergasser *et al.* 2012) to PCR amplify each putative SNP together with 100–200 bp of flanking sequence. Each locus was PCR amplified in one fur seal individual that had been genotyped as homozygous at that locus and one that had been genotyped as heterozygous. PCRs was carried out using 1.5 μL of template DNA, 20 mM Tris–HCl (pH 8.3), 100 mM KCl, 2 mM MgCl2, 10x Reactionbuffer Y (Peqlab), 0.25 mM dNTPs, 0.25 mol/L of each primer, and 0.5U of Taq DNA polymerase (VWR). The following PCR profile was used: one cycle of 1.5 min at between 59° and 62° depending on the primers used (Supplementary Table 2), 60 sec at 72°C; and one final cycle of 7 min at 72°C. 5 μL of the resulting PCR product was then purified using shrimp alkaline phosphatase and exonuclease I (NEB) following the manufacturer’s recommended protocol. All fragments were then sequenced in both directions using the Applied Biosystems BigDye Terminator v3.1 Cycle Sequencing Kit (Thermo Fisher Scientific) and analyzed on an ABI 3730xl capillary sequencer. Forward and reverse reads were aligned using Geneious v10.2.3 (Kearse *et al.* 2012). Heterozygous sites were identified as those with two peaks of roughly equal intensity but with around half the intensity of a homozygote.

### Linkage disequilibrium decay

Prior to quantifying linkage disequilibrium, we filtered the SNP dataset as shown in Figure S1A. First, to minimise the occurrence of unreliable genotypes, we removed individual genotypes with a depth of coverage below eight or above 30 using VCFtools (Danecek *et al.* 2011). Genotypes with very low depth of coverage have a greater likelihood of being called incorrectly as it can be difficult to distinguish between homozygotes and heterozygotes when very few reads are present. Similarly, genotypes with very high depth of coverage are more likely to be spurious as high coverage can result from misalignment due to the presence of paralogous loci or repeats (Fountain *et al.* 2016). Second, as including SNPs from short scaffolds can downwardly bias LD values, we retained only SNPs located on the longest 100 scaffolds of the assembly (min length = 6.6 Mb, max length = 33.1 Mb). Third, as an additional quality filtering step, we used information on known parental relationships to identify loci with Mendelian incompatibilities using the mendel function in PLINK v1.9 and removed these from the dataset. Fourth, to remove any possible confounding effects of population structure, we focussed on the single largest population of South Georgia. Fifth, to provide an informative dataset while further minimising genotyping error, we discarded SNPs with a MAF of less than 0.1 and/or called in less than 50% of individuals using PLINK. As a final quality control step, we also removed SNPs that did not conform to Hardy-Weinberg equilibrium (HWE) with a *p*-value threshold < 0.001 using the --hwe function in PLINK.

Using the final dataset of 27,347 SNPs genotyped in 57 individuals (Figure S1A), we used the --r2 function in PLINK to quantify pairwise LD between all pairs of SNPs located within 500 kb of each other. We visualised LD decay with distance by fitting a nonlinear regression curve using the nls package in R, where the expected value of *r*^2^ under drift-recombination (*E*(*r*^2^)) was expressed according to the Hill and Weir function (Hill and Weir 1988), as implemented by Marroni *et al.* (2011):

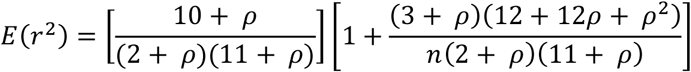

where *N*_*e*_ is the effective population size, *c* is the recombination fraction between sites, *p* = *4N*_*e*_*c* and *n* is the number of scaffolds (Remington *et al.* 2001).

### Population structure

Prior to quantifying population structure, we filtered the full SNP dataset as shown in Figure S1B. We did not initially filter the dataset for SNPs with low depth of coverage as for the analysis of population structure we wanted to retain as many SNPs as possible that were genotyped across all the populations. We also did not remove individuals with high levels of missing data in order to maximise the representation of all populations in the final dataset. Nevertheless, because closely related individuals can bias population genetic structure analysis by introducing both Hardy-Weinberg and linkage disequilibrium (Rodriguez-Ramilo and Wang 2012; Wang 2017), we used known parentage information to remove adults and related pups (full and half siblings) from the South Georgia dataset. Second, SNPs with a MAF of less than 0.05 and/or called in less than 99% of individuals were discarded using VCFtools. Third, SNPs were pruned for LD using the --indep function in PLINK. We used a sliding window of 50 SNPs, a step size of 5 SNPs and removed all variants in a window above a variance inflation factor threshold of 2, corresponding to *r*^2^ = 0.5. As population structure can lead to deviations from HWE, we did not filter our final dataset for HWE.

Using the final dataset of 28,062 SNPs genotyped in 37 individuals (Figure S1B), we first visualised population structure by performing a principal components analysis (PCA) using the R package adegenet (Jombart 2008). We then used a Bayesian clustering algorithm implemented by the program STRUCTURE to identify the number of genetic clusters (*K*) present in the dataset. We performed STRUCTURE runs for values of *K* ranging from one to six, with five simulations for each *K* and a burn-in of 100,000 iterations followed by 1,000,000 Markov chain Monte Carlo iterations. We used the admixture and correlated allele frequency models without sampling location information. The R package pophelper (Francis 2017) was then used to analyse the STRUCTURE results, parse the output to CLUMPP for averaging across iterations and for visualising individual assignment probabilities. The optimal *K* was selected based on the maximum value of the mean estimated *ln* probability of the data (Ln Pr(*X* | *K*) as proposed by Pritchard *et al.* (2000) and the Δ*K* method of Evanno *et al.* (2005). For comparison, we also implemented the above analyses using microsatellite data for the same individuals.

### Inbreeding coefficients

Prior to quantifying inbreeding, we filtered the SNP dataset as shown in Figure S1C. First, for the analysis of inbreeding we wanted a dataset with as few gaps as possible so we discarded one individual with more than 90% missing data. Second, we removed individual genotypes with a depth of coverage below eight or above 30 using vcftools. Third, we removed loci with Mendelian incompatibilities, and fourth, we again restricted the dataset to the focal population of South Georgia. Fifth, we discarded SNPs with a MAF of less than 0.05 and/or called in less than 75% of individuals using vcftools. Finally, we filtered the SNPs for HWE as described previously and pruned linked SNPs out of the dataset using the --indep function in PLINK with the parameters shown above.

Using the final dataset of 9,853 SNPs genotyped in 56 individuals (Figure S1C), we calculated four genomic estimates of individual inbreeding: standardised multi-locus heterozygosity (sMLH), an estimate based on the variance of additive genotype values 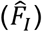, an estimate based on excess homozygosity 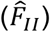 and an estimate based on the correlation of uniting gametes, which gives more weight to homozygotes of the rare allele at each locus 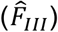. The former was calculated using the sMLH function in the R package inbreedR (Stoffel *et al.* 2016) whilst the latter were calculated in GCTA v1.24.3 (Yang *et al.* 2011). To test for a significant correlation in heterozygosity across marker loci, we quantified identity disequilibrium (ID) using the measure *g*_2_ in the R package inbreedR (Stoffel *et al.* 2016) where significant *g*_2_ values provide support for variance in inbreeding in the population. Finally, we compared the resulting *g*_2_ value with the variance in our inbreeding coefficients to determine the expected correlation between estimated 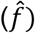 and realized (*f*\*) level of inbreeding (Szulkin *et al.* 2010) given as:

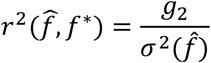

## RESULTS

### Hybrid genome assembly

We used PacBio sequencing to improve an existing Antarctic fur seal genome assembly. Using PBJelly, we were able to close a total of 45,394 gaps, resulting in a 40% reduction in overall gap space (assembly v1.2, Table 1). Subsequent assembly correction with Quiver resulted in a total of 11,319,546 modifications to the PBJelly assembly consisting of 291,179 insertions, 1,117,226 substitutions and 9,911,141 deletions. Finally, PILON corrected 653,246 homozygous insertions (885,794 bp), 87,818 deletions (127,024 bp) and 34,438 single-base substitutions and closed an additional 2,170 gaps in the Quiver assembly. Overall, gap closing and error correction resulted in a hybrid Antarctic fur seal assembly with a total length of 2.3 Gb (v1.4, Table 1). The number of scaffolds in the genome was reduced from 8,126 to 6,169 such that 50% of the final assembly is now contained within the longest 108 scaffolds (Table 1).

### Genome synteny

To investigate synteny between the Antarctic fur seal and the dog, we aligned the fur seal scaffolds to the dog genome (CanFam3.1). We estimated overall sequence divergence between the two species to be 13.8%. Visualisation of the full alignment revealed that all of the dog chromosomes are represented in the fur seal assembly (Figure S2). Alignment of the 40 longest fur seal scaffolds (min length = 10.7 Mb, max length = 33.1 Mb) revealed strong chromosomal synteny between the two genomes, with the vast majority of the fur seal scaffolds mapping exclusively or mainly to a given dog chromosome (Figure 2). Specifically, for 37 of the scaffolds, over 90% of the total alignment length was to a single dog chromosome, with 26 of those aligning exclusively to a single dog chromosome. Only one scaffold (S4 in Figure 2) aligned in roughly equal portions to two different dog chromosomes (62% to D5 and 38% to D26).

**Table 1.** Genome assembly statistics for successive improvements of the original Antarctic fur seal genome assembly.

|  | **v1.0.2** | **v1.1** | **v1.2** | **v1.3** | **v1.4** |
| --- | --- | --- | --- | --- | --- |
|  | **ALLPATHS3** | **GapCloser** | **PBJelly2** | **Quiver** | **Pilon** |
| Number of scaffolds | 8,126 | 8,126 | 6,170 | 6,170 | 6,169<sup>†</sup> |
| N90<sup>a</sup> | 890,836 (768) | 888,912 (768) | 1,624,547 (387) | 1,511,352 (387) | 1,542,705 (387) |
| N50<sup>a</sup> | 3,169,165 (233) | 3,165,747 (233) | 6,454,664 (108) | 6,076,522 (108) | 6,207,322 (108) |
| N10<sup>a</sup> | 8,459,351 (25) | 8,458,289 (25) | 17,733,103 (11) | 16,529,571 (11) | 16,861,656 (11) |
| Longest scaffold (bp) | 13,012,173 | 12,999,316 | 34,690,325 | 32,399,786 | 33,062,611 |
| Total size (bp) | 2,405,038,055 | 2,403,626,805 | 2,426,014,533 | 2,268,217,244 | 2,313,485,084 |
| Gaps present (%) | 4.79 | 3.26 | 0.62 | 0.57 | 0.55 |
| Number of gaps | 136,284 | 90,432 | 45,102 | 22,783 | 20,613 |
| Average gap size (bp) | 845.56 | 866.87 | 331.16 | 570.37 | 613.39 |

<sup>a</sup> Size in bp (number of scaffolds)
<sup>†</sup> Excluding the mitochondrial genome, which was filtered out by Pilon

**Figure 2.**
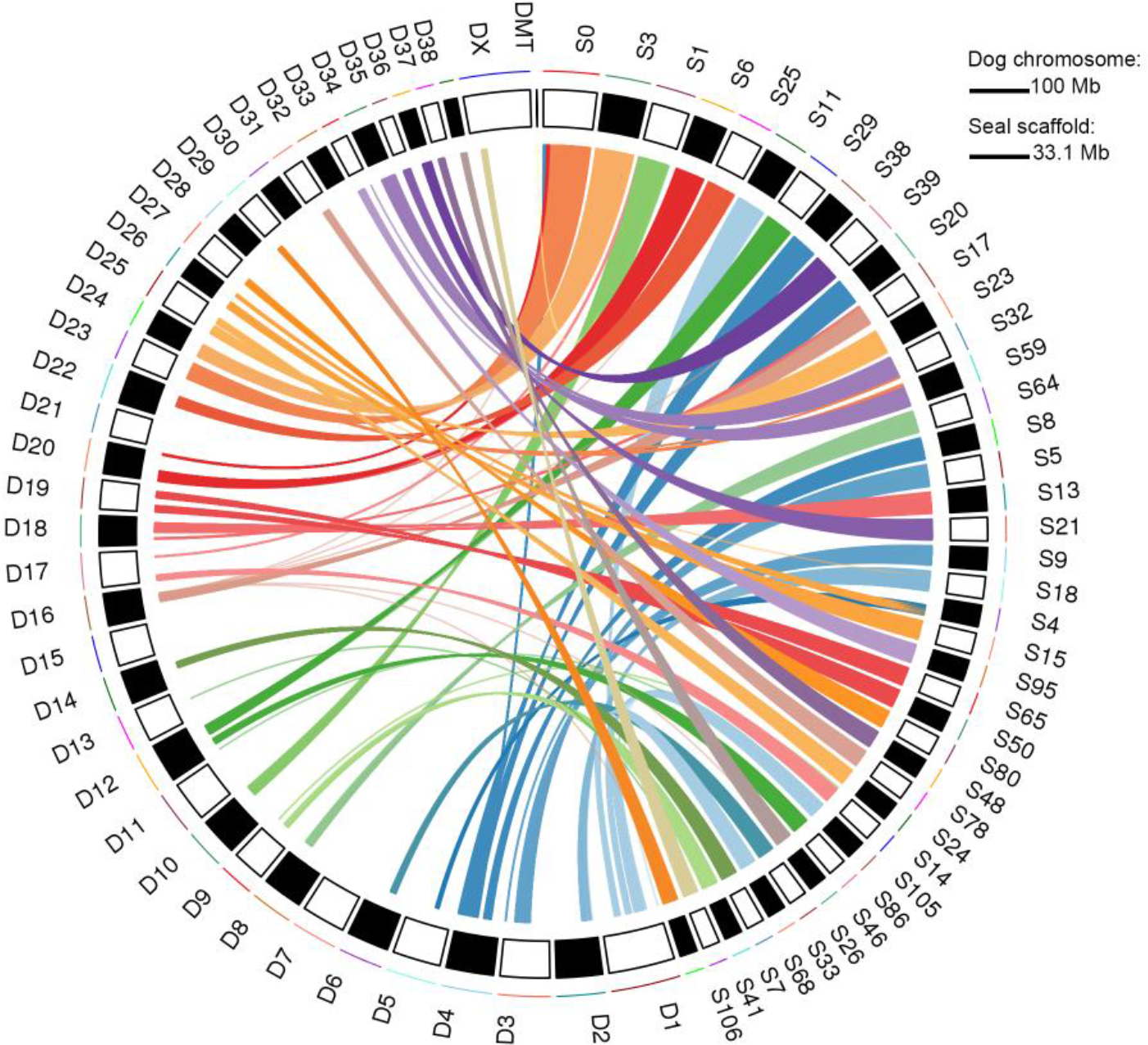
Synteny of the longest 40 Antarctic fur seal scaffolds (10.7-33.1 Mb; right, prefixed S) with dog chromosomes (left, prefixed D). Mapping each fur seal scaffold to the dog genome resulted in multiple alignment blocks (mean = 2.1 kb, range = 0.1-52.8 kb) and alignments over 5 kb are shown.

### RAD sequencing and SNP discovery

RAD sequencing of 83 fur seal individuals generated an average of 5,689,065 250bp paired-end reads per individual. After mapping these reads to the reference genome, a total of 677,607 biallelic SNPs were discovered using GATK’s best practices workflow for variation discovery (see Materials and methods for details), with an average coverage of 727. We then filtered the dataset in three different ways (Supplementary Figure 1) to generate datasets suitable for the analysis of LD decay, population structure and inbreeding.

### SNP validation

To provide an indication of the quality of our SNP dataset, we used Sanger sequencing to validate 50 randomly selected loci. For each locus, we sequenced a single heterozygote and a single homozygote individual based on the corresponding GATK genotypes. For 40 of these loci, we successfully obtained genotypes for both individuals (Supplementary Table 2). Concordance between the GATK and Sanger genotypes was high, with 76 / 80 genotypes being called identically using both methods, equivalent to a validation rate of 95%. The four discordant genotypes were all initially called as homozygous with GATK but subsequently validated as heterozygous with Sanger sequencing.

### LD decay

The pattern of LD decay within South Georgia was quantified based on 27,347 SNPs genotyped in 57 individuals and located on the 100 longest fur seal scaffolds. LD was found to decay rather rapidly, with *r*^2^ reaching the background level (average *r*^2^ = 0.12) by around 18 kb and decreasing to values approaching zero by around 350 kb (Figure 3). Strong LD (*r*^2^ >= 0.5) decayed by around 5 kb and moderate LD (*r*2 >= 0.2) by around 7 kb.

**Figure 3.**
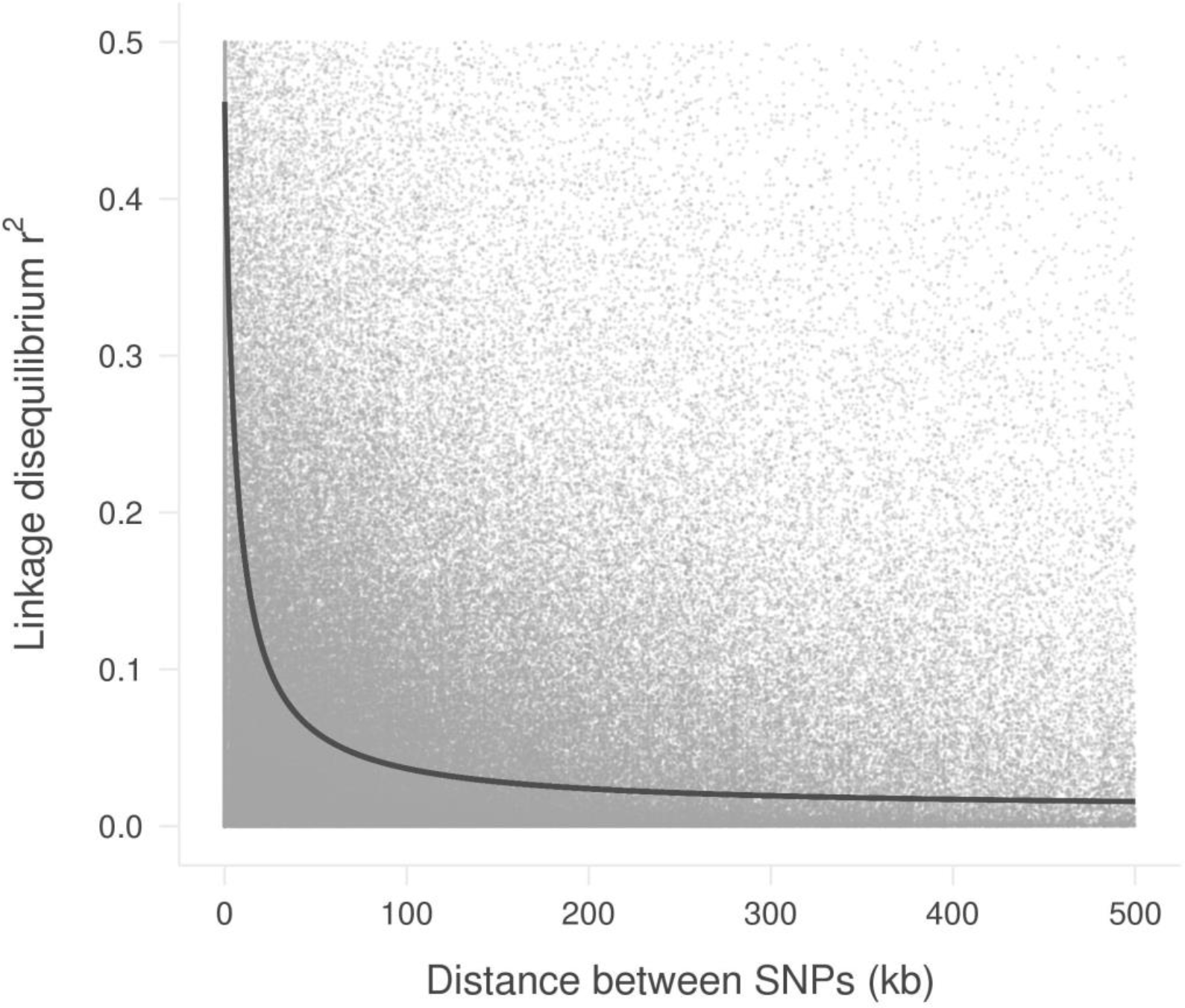
Plot of linkage disequilibrium (r^2^) against distance between SNPs in the Antarctic fur seal. LD was calculated using 27,347 filtered RAD SNPs from the 100 largest scaffolds of 57 South Georgia individuals. Grey dots indicate observed pairwise LD. Dark grey curve shows the expected decay of LD in the data estimated by nonlinear regression of r^2^.

### Population structure

Finally, we used a dataset of 37 pups genotyped at 27 microsatellites and 28,062 SNPs to quantify the pattern and strength of population structure across the species’ circumpolar range. PCA of the microsatellite dataset uncovered weak clustering with South Georgia, the South Shetlands and Bouvetøya tending to separate apart from Kerguelen, Heard and Macquarie Islands along the first PC axis (Figure 4A). However, considerable scatter and no clear pattern of separation was found along either PC2 or PC3 (Figures 4A and 4C). By contrast, population structure was more clearly defined in the PCA of the SNP dataset. Specifically, the first PC axis clearly resolved two distinct island groups, the first comprising South Georgia, the South Shetlands and Bouvetøya and the second comprising Kerguelen, Heard Island and Macquarie Island (Figure 4B). Within the first island group, Bouvetøya clustered apart from South Georgia and the South Shetlands along PC2 (Figure 4B) while all three locations clustered apart from one another along PC3 (Figure 4D).

To test whether population structure could be detected without prior knowledge of the sampling locations of individuals, we used a Bayesian approach implemented within STRUCTURE (Pritchard *et al.* 2000). This program works by partitioning the data set in such a way that departures from Hardy-Weinberg and linkage equilibrium within the resulting groups are minimized. Separately for the microsatellite and SNP datasets, five replicate runs were conducted for each possible number of groups (*K*) ranging from one, implying no population differentiation, through to six, which would imply that all of the populations are genetically distinct. For the microsatellite dataset, Ln Pr(*X* | *K*) and Δ*K* both peaked at 2, indicating support for the presence of two genetically distinct populations (Figure S3A and C). Membership coefficients for the inferred groups are summarized in Figure S4A and indicate the presence of a Western population comprising individuals from South Georgia, the South Shetlands and Bouvetøya, and an Eastern population comprising individuals from Kerguelen, Heard Island and Macquarie Island.

**Figure 4.**
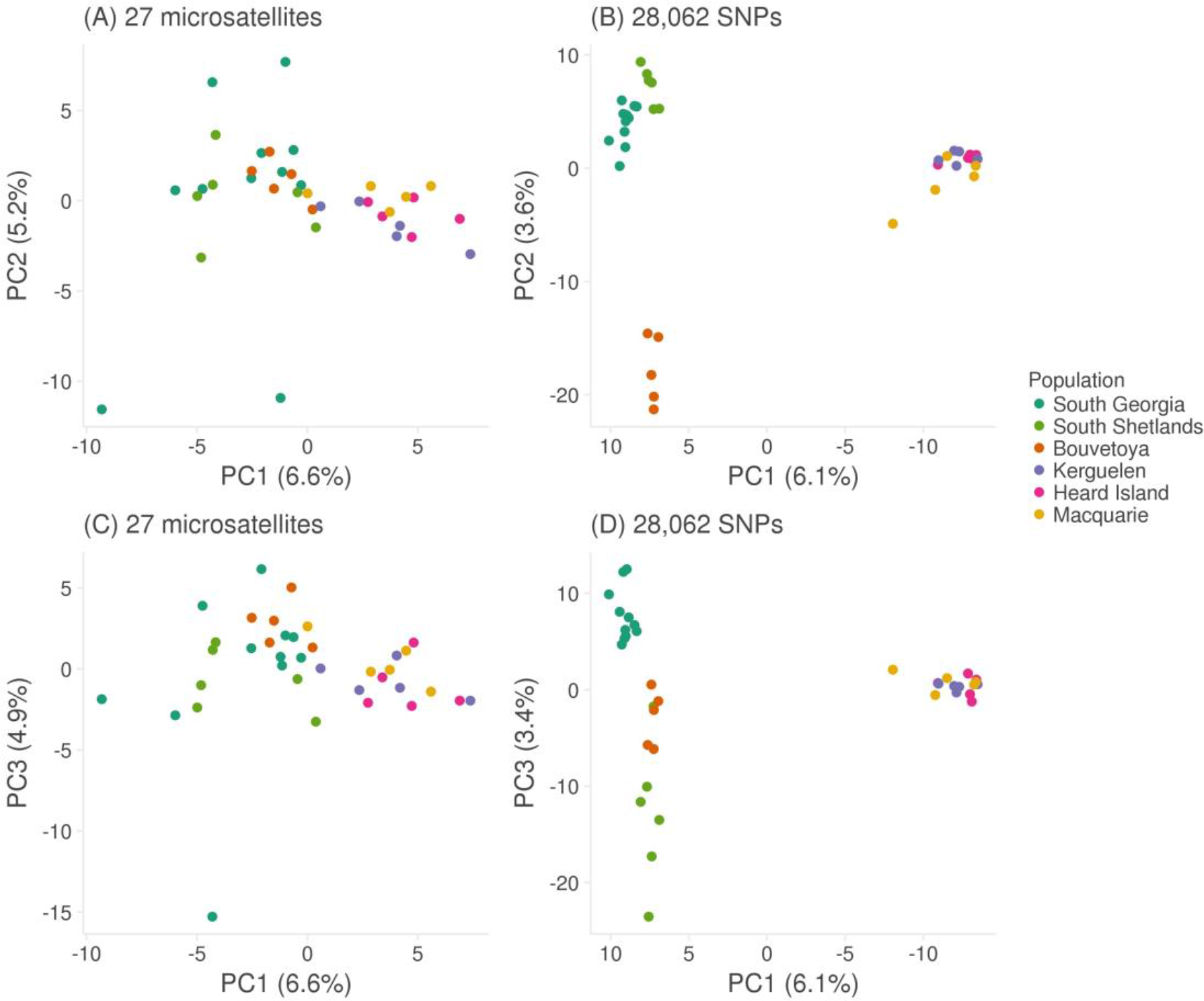
Scatterplots showing individual variation in principal components (PCs) one and two (A and B), and one and three (C and D) derived from a principal component analysis conducted using 27 microsatellites (A and C) and 28,062 SNPs (B and D). Variance explained by each PC is shown in brackets.

For the RAD dataset, Ln Pr(*X* | *K*) also peaked at 2 but remained high for *K* = 3 and 4, while Δ*K* reached its maximum at *K* = 4 (Figure S3B and D). To explore this further, we plotted membership coefficients for *K* = 2 to 6 for both the microsatellite and SNP datasets. For the former, no evidence of population structure was found beyond *K* = 2, with successive increases in *K* merely introducing additional admixture (Figure S4A). By contrast for the latter, plots corresponding to *K* values greater than 2 clearly resolved further hierarchical structure (Figure S4B). Results for *K* = 4 are shown in Figure 1, in which Kerguelen, Heard and Macquarie Islands are resolved as a single population, while South Georgia, the South Shetlands and Bouvetøya can be readily distinguished based on their corresponding group membership coefficients.

### Inbreeding

Inbreeding in the focal population at South Georgia was investigated using data from 9,853 SNPs genotyped in 56 individuals (Figure 5A). Identity disequilibrium differed significantly from zero (0.0052; bootstrap 95% confidence interval = 0.0008-0.0091, *p* = 0.023, Figure 5B) providing evidence for variance in inbreeding within the sample of individuals. Each individual’s level of inbreeding was quantified from the SNP dataset using four different genomic inbreeding coefficients (sMLH, 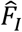, 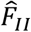 and 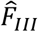, see Materials and methods for details). All four of these measures were inter-correlated, with correlation coefficients (*r*) ranging from 0.69 to 0.83. (Figure 5C-E). Furthermore, the variances of 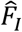, 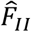 and 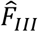 fell within the 95% confidence interval of *g*_2_, suggesting that the expected correlation between the estimated and realized level of inbreeding does not differ significantly from one (Figure 5B).

**Figure 5.**
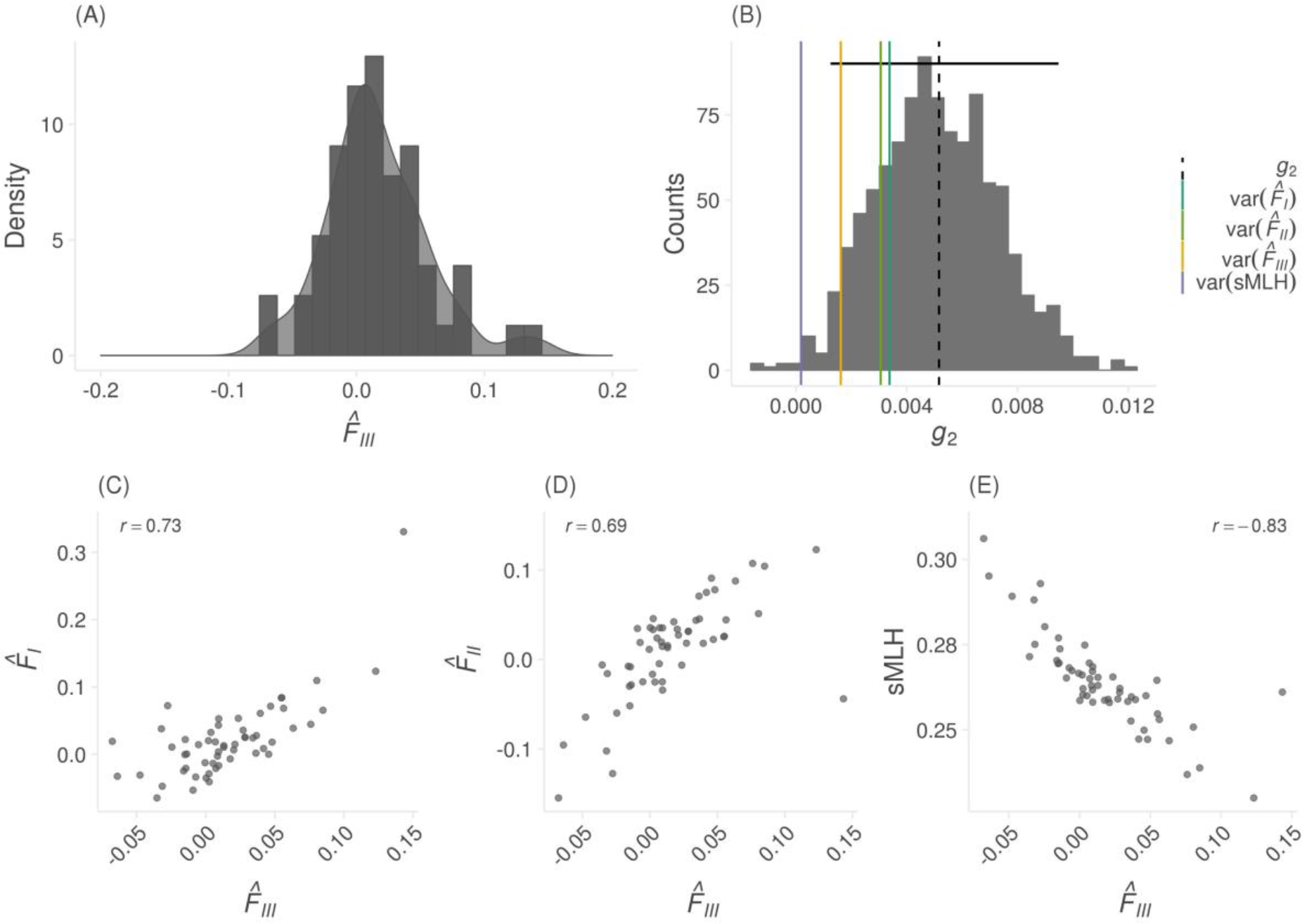
Distribution of inbreeding coefficients 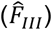 for 56 South Georgia individuals (A). Distribution of identity disequilibrium (*g*_2_) estimates from bootstrapping over individuals (B). Horizontal black line shows 95% confidence interval from 1000 bootstrap replications. Vertical dashed line represents empirical *g*_2_ estimate. Vertical coloured lines represent variance in inbreeding coefficients. Pairwise correlation between 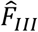 and 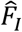, 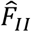, and sMLH based on 9,853 SNPs (C, D, E). Pearson’s correlation coefficients are shown.

## DISCUSSION

Advances in high throughput sequencing technology have afforded researchers the opportunity to generate genome assemblies and genomic marker datasets for virtually any species for which high quality DNA can be collected. These resources allow a broad range of questions in ecology and evolution to be addressed with greater power and precision than was possible with traditional methods. In this study, we utilised PacBio sequencing to refine an existing Antarctic fur seal genome assembly and combined this with RAD sequencing to characterize synteny with the dog genome, elucidate the rate of LD decay, resolve global population structure and quantify the variance in inbreeding. Our results provide new insights at multiple levels of organisation that enrich our understanding of an important Antarctic marine top predator and indicate the general promise of these and related approaches for tackling broad-reaching questions in population and evolutionary genetics.

### Genome alignment

An important outcome of this study is a significantly improved Antarctic fur seal genome assembly. This was achieved through three iterative steps involving gap filling, inclusion of long PacBio reads and assembly error correction respectively. Overall, the number of scaffolds was reduced by around one quarter, while N50 almost doubled to over 6 Mb and the proportion of gaps was reduced by around an order of magnitude to around half a percent. This represents an improvement over existing pinniped assemblies such as the walrus (*Odobenus rosmarus divergens*, GenBank accession number GCA_000321225.1) and Weddell seal (*Leptonychotes weddellii*, GenBank accession number GCA_000349705.1), which both have lower N50 values (2.6 and 0.9 Mb respectively). The improved Antarctic fur seal genome will therefore serve as an important resource for the wider pinniped community. However, there is still considerable room for improvement as a handful of other marine mammal genome assemblies incorporating longer range information show higher levels of contiguity e.g. killer whale, (*Orcinus orca*, N50 = 12.7 Mb) and Hawaiian monk seal (*Neomonachus schauinslandi*, N50 = 22.2 Mb) (Foote *et al.* 2015; Mohr *et al.* 2017).

To further quantify genome quality and to explore patterns of synteny, we mapped the scaffolds of our new assembly to the dog genome. The resulting alignment revealed almost complete coverage of the dog chromosomes. This is in line with the observation that the total length of the assembly has not changed appreciably between versions and suggests that the assembly is near-complete, with the exception of the Y-chromosome for which sequence data are currently lacking as the genome individual is a female. In general, carnivore genomes show high levels of synteny (Arnason 1974; Ferguson-Smith and Trifonov 2007), with pinnipeds in particular exhibiting highly conserved karyotypes indicative of slow rates of chromosomal evolution (Beklemisheva *et al.* 2016). By contrast, the domestic dog has an extensively re-arranged karyotype differentiated from the ancestral carnivore karyotype by over 40 separate fission events (Nie *et al.* 2011). To provide insights into the extent of conservation of chromosomal blocks between seals and dogs, we mapped the longest 40 fur seal scaffolds to the dog genome. We found a clear pattern whereby all but one of the scaffolds mapped exclusively or mainly to single chromosomes, indicating the conservation of large genomic tracts often several Mb in length. The remaining scaffold mapped to two dog chromosomes in roughly equal proportions, suggestive of either a fission event in the lineage leading to dogs or a fusion event in the lineage leading to seals. By focusing only on the largest scaffolds, we had little power to detect multiple chromosomal rearrangements, although these are to be expected given a substantial increase in the number of chromosomes in dogs (2n = 74) relative to the seal (2n = 36) (Gustavsson 1964; Arnason 1974)). Nevertheless, the observed high degree of synteny is consistent with previous studies revealing both strong sequence homology and the conservation of polymorphic loci between seals and dogs (Osborne *et al.* 2011; Hoffman *et al.* 2013).

### SNP discovery and validation

Our study found a total of 667,607 SNPs in a discovery pool of 83 individuals. These markers will be useful for future studies including the planned development of a high-density SNP array. However, not all SNPs are suitable for every analysis due to differential sensitivity to missing data, low depth of sequencing coverage and the inclusion of low frequency alleles (Shafer *et al.* 2017). Similarly, filtering for deviations from HWE and Mendelian incompatibilities should reduce the error rate by reducing the frequency of erroneous genotypes. Yet, as population structure can generate deviations from HWE, stringent filtering may also remove genuine signal. We therefore carefully considered how best to filter our SNP dataset for each of our main analyses. For LD decay, we applied relatively strict filters as we sought a high-quality dataset with consistently high coverage across individuals. For population structure, it was important to have as many SNPs as possible represented in all of the sampling locations, so we did not remove genotypes with low coverage or containing Mendelian incompatibilities but instead filtered to retain SNPs genotyped in at least 99% of individuals. Conversely, for the estimation of inbreeding, we honed in on a reduced subset of higher quality SNPs with greater average depth of coverage, in Hardy-Weinberg and linkage equilibrium, and with no evidence of Mendelian incompatibilities.

Even with stringent filtering, it is possible to retain SNPs in a dataset that have been called incorrectly. We therefore attempted to validate 50 randomly selected loci by Sanger
sequencing selected individuals with homozygous and heterozygous genotypes as determined from the RAD data. For the 40 loci that we were able to successfully sequence, around 95% of the Sanger genotypes were identical to the RAD genotypes. Although this validation step required additional experimental effort, our results compare favourably with other studies (Cruz *et al.* 2017; Bourgeois *et al.* 2018) and thus give us confidence in the overall quality of our data.

### Linkage disequilibrium decay

We used the genomic positions of SNPs mapping to the largest 100 scaffolds to quantify the pattern of LD decay in the focal population of South Georgia. We found that LD decays rapidly, with moderate LD extending less than 10 kb. This is despite the species having experienced a population bottleneck in the 19^th^ century which is expected to increase LD. A direct comparison with other organisms is hindered both by a paucity of data for most species and by the use of different measures for quantifying LD. However, our results are broadly in line with other wild vertebrate populations such as polar bears, Alaskan gray wolves and flycatchers, where moderate LD also extends less than 10 kb (Gray *et al.* 2009; Malenfant *et al.* 2015; Kardos, Husby, *et al.* 2016). Extended LD has been documented in a number of species but in most cases this is associated with extreme bottlenecks, such as those experienced during domestication (Harmegnies *et al.* 2006; McKay *et al.* 2007; Meadows *et al.* 2008). Although Antarctic fur seals are generally believed to have also experienced a very strong historical bottleneck, a recent Bayesian analysis suggested that this may have been less severe than thought, with the effective population size probably falling to several hundred (Hoffman *et al.* 2011). Furthermore, the population recovered from the bottleneck within a few generations, which could have mitigated the increased genetic drift and inbreeding effects that elevate and maintain strong LD. Additionally, the population is currently estimated to number around 2–3 million individuals (Boyd 1993) and is one of the most genetically diverse pinnipeds (Stoffel *et al.* unpublished results). Therefore, given that LD is a function of both recombination rate and population size (Hill 1981), the rapid decay of LD in this species might also be a reflection of high long-term effective population sizes.

### Population structure

To provide further insights into the recovery of Antarctic fur seals globally, we quantified population structure across the species’ geographic range. Microsatellite genotypes provided evidence for two major geographic clusters, the first corresponding to South Georgia, the South Shetlands and Bøuvetoya, and the second corresponding to Kerguelen, Heard and Macquarie Island. By contrast, the RAD data uncovered an additional level of hierarchical structure, resolving South Georgia, the South Shetlands and Bøuvetoya as distinct populations. This is consistent with simulation studies suggesting that thousands of SNPs should outperform small panels of microsatellites at resolving population structure (Haasl and Payseur 2011) as well as with more recent empirical studies that have directly compared microsatellites with SNPs (Rašić *et al.* 2014; Vendrami *et al.* 2017). Furthermore, many of our populations had sample sizes of around five individuals yet could still be clearly distinguished from one another. This is in line with a recent simulation study suggesting that sample sizes as small as four individuals may be adequate for resolving population structure when the number of markers is large (Willing *et al.* 2012). Thus, our results have positive implications for studies of threatened species for which extensive sampling can be difficult but where understanding broad as well as fine-scale population structure is of critical importance.

It is generally believed that Antarctic fur seals were historically extirpated from virtually all of their contemporary breeding sites across the sub-Antarctic, with the possible exception of Bøuvetoya, where sealing expeditions were more sporadic (Christensen 1935) and around a thousand breeding individuals were sighted just a few decades after the cessation of hunting (Olstad 1928). South Georgia was the first population to stage a major recovery, probably because a number of individuals survived at isolated locations inaccessible to sealers around the South Georgia mainland (Bonner 1968). Consequently, several authors have speculated that emigrant individuals from the expanding South Georgia population may have recolonized the species former range (Boyd 1993; Hucke-Gaete *et al.* 2004). However, Wynen *et al.* (2000) resolved two main island groups with mtDNA, while Bonin *et al.* (2014) found that significant differences between the South Shetland Islands and South Georgia with microsatellites. Our results build on these studies in two ways. First, the two major clusters we resolved using both microsatellites and RAD sequencing are identical to those identified by Wynen *et al.* (2000), suggesting that broad-scale population structure is not simply driven by female philopatry but is also present in the nuclear genome. Second, within the Western part of the species range, we not only found support for the South Shetlands being different from South Georgia, but also Bøuvetoya, suggesting that relict populations probably survived at all three of these locations. By contrast, no sub-structure could be detected within the Eastern part of the species range, which taken at face value might suggest that a single population survived sealing in this region. Consistent with this, historical records suggest that fur seals went locally extinct at Heard and Macquarie islands (Page *et al.* 2003; Goldsworthy *et al.* 2009) and these populations may therefore have been recolonised by surviving populations in the Kerguelen archipelago. Thus, our study highlights the importance of relict populations to species recovery while also providing some evidence for local extinctions having occurred.

### Inbreeding

Delving a level deeper, we investigated individual variation in the form of inbreeding. A recent meta-analysis has shown that small panels of microsatellites are almost always underpowered to detect variation in inbreeding (Szulkin *et al.* 2010; Miller *et al.* 2013). By contrast, a handful of recent studies have shown that tens of thousands of SNPs are capable of accurately quantifying inbreeding (Hoffman *et al.* 2014; Huisman *et al.* 2016; Berenos *et al.* 2016; Chen *et al.* 2016; Kardos *et al.* 2018). While empirical studies to date have largely focused on small, isolated populations where inbreeding may be common, it is less clear how prevalent inbreeding could be in larger, free-ranging populations. We found several lines of evidence in support of inbreeding in fur seals. First, *g*_2_ was significantly positive indicating identity disequilibrium within the samplel of individuals. Second, the variance of the genomic 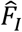, 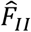 and 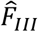 were found to lie within the 95% confidence intervals of *g*_2_ and therefore we can expect our estimates to reflect the realized level of inbreeding in the population. Third, the genomic inbreeding coefficients were strongly inter-correlated, suggesting that our markers are uncovering consistent information about variation in genome-wide homozygosity caused by inbreeding.

Our results are surprising given that Antarctic fur seals number in the millions and are free-ranging and highly vagile. However, the species is also highly polygynous, with a handful of top males fathering the majority of offspring (Hoffman *et al.* 2004) and females exhibiting strong natal site fidelity (Hoffman and Forcada 2012) which could potentially lead to matings between close relatives. As demographic effects can also generate variance in inbreeding *sensu lato*, we also cannot discount the possibility that the historical bottleneck contributed towards the variation we see today. To test this, we would need to quantify the length distribution of runs of homozygosity, which would require denser SNP data.

Our work builds upon another recent study that used RAD sequencing to quantify inbreeding in wild harbour seals (Hoffman *et al.* 2014) where a higher estimate of *g*_2_ was found, indicative of a greater variance in inbreeding within the sample. However, the study focused on stranded seals, many of which died of lungworm infection and may therefore have been enriched for unusually inbred individuals. In the current study, pups were sampled at random from within a single breeding colony, together with their parents. Consequently, our sample should be
more representative of the underlying distribution of inbreeding within the population. In line with this, our estimate of *g*_2_ is more similar to those obtained in wild populations of other polygynous mammals such as Soay sheep and red deer (Huisman *et al.* 2016; Berenos *et al.* 2016).

Our results are consistent with previous studies documenting HFCs for numerous traits in the South Georgia population (Hoffman *et al.* 2004; 2010; Forcada and Hoffman 2014) and suggest that these may well reflect inbreeding depression. More generally, literally hundreds of studies have documented HFCs across the animal kingdom (Coltman and Slate 2003) and it has been strongly argued that these HFCs are highly unlikely to occur when there is no variance in inbreeding (Szulkin *et al.* 2010). The fact that we found variation in inbreeding in a large, free-ranging population is consistent with this notion and therefore contributes towards a growing body of evidence suggesting collectively that inbreeding could be more common in wild populations than previously thought.

### Conclusion

We have generated an improved genome assembly for an important Antarctic marine top predator and used RAD sequencing to provide diverse insights from the level of the species through the population to the individual. Focusing on the larger South Georgia population, we characterised rapid LD decay and uncovered significant variation in individual inbreeding, while population-level analyses resolved clear differences among island groups that emphasise the importance of relict populations to species recovery. RAD sequencing and related approaches might conceivably be applied to other wild species to characterise patterns of LD decay, elucidate fine scale population structure and uncover the broader prevalence of inbreeding and its importance to wild populations.

## ACKNOWLEDGEMENTS

We thank the British Antarctic Survey’s field assistants on Bird Island for the collection of tissue samples and to David Vendrami and Martin Stoffel for useful discussions about SNP filtering. We would also like to acknowledge Anika Winkler who carried out the Illumina sequencing. Samples from Cape Shirreff, South Shetland Islands were collected under U.S. Marine Mammal Protection Act permit #16074. This work contributes to the Ecosystems project of the British Antarctic Survey, Natural Environmental Research Council, and is part of the Polar Science for Planet Earth Programme. This project was funded by a Deutsche Forschungsgemeinschaft standard grant (HO 5122/3-1) and also supported by the Swedish Research Council FORMAS (231-2012-450) as well as core funding from the Natural Environment Research Council to the British Antarctic Survey’s Ecosystems Program. KKD was supported by the Natural Environment Research Council (grant NE/K012886/1) and AMB was supported by the Knut and Alice Wallenberg Foundation as part of the National Bioinformatics Infrastructure Sweden at SciLifeLab.

## AUTHOR CONTRIBUTIONS

EH and JIH conceived and designed the study. IG and JIH carried out the DNA extractions and microsatellite genotyping. KKD carried out the RAD library preparation. A-CP performed the SNP validation. JF, SG, MG, KKD, JK, JIH and JW contributed materials and funding. AMB assembled the new version of the genome with input from JW. EH carried out the SNP calling and analysed the data. EH and JIH wrote the first version of the manuscript. All of the authors commented on and approved the final manuscript.

